# EnrichMiner: a biologist-oriented web server for mining biological insights from functional enrichment analysis results

**DOI:** 10.1101/2023.07.12.548786

**Authors:** Xian Liu, Kaikun Xu, Xin Tao, Xiaochen Bo, Cheng Chang

**Author notes:** To whom correspondence should be addressed: Xian Liu, Xiaochen Bo, Cheng Chang. These authors contributed equally: Xian Liu and Kaikun Xu.

## Abstract

Functional enrichment analysis has been widely used to help researchers obtain biological insights from -omics data. However, the results are often redundant and difficult to digest. The key is developing tools to help users explore the relationships between the enriched terms, remove the redundant terms, and finally select representative terms. However, existing tools hardly make a good integration between enrichment analysis and representative terms selection in a biological-friendly manner. Here, we developed a biologist-oriented web server named EnrichMiner to provide a one-stop solution. It is a complete analysis pipeline from a gene list or a ranked gene table to published-style figures. More importantly, it provides user-friendly interfaces and rich interactive operations to help users explore the term relationships and remove redundancy. EnrichMiner has been integrated into the ExpressVis platform, and is freely accessible at https://omicsmining.ncpsb.org.cn/ExpressVis/EnrichMiner and does not require login.

## 1 Introduction

Omics technologies, such as RNA-seq and mass spectrometry, have been widely used in biomedical research. One of the main analysis methods to mine biological insights from these omics data is functional enrichment analysis. At present, the most commonly used enrichment analysis methods are Over Representation Analysis (ORA)^1^ and Gene Set Enrichment Analysis (GSEA)^2^. ORA takes a gene/protein list of interest as input and iteratively tests whether one gene set (such as all the genes in a pathway) contains disproportionate genes of the input. Differently, GSEA takes a gene table with ranked metrics as input and tests whether genes of a gene set accumulate at the top or bottom of the full gene vector ordered by the quantitative values^3^. Many excellent tools have been developed for the two enrichment methods, such as DAVID^4^, g:Profiler^5^, Enrichr^6^ and WebGestalt^7^. Meanwhile, similar analysis methods such as Metabolite Set Enrichment Analysis (MSEA)^8^ and Kinase–Substrate Enrichment Analysis (KSEA)^9^ have also been developed to identify the significant metabolites or kinase’s activity.

However, digesting the enriched terms remains a big challenge. The background gene set library usually contains large number of gene sets with overlapping genes and the ORA and GSEA methods consider each term independently. Thus, they usually output the enriched terms with high redundancy. Some tools have been developed to remove the redundancy of enriched terms to make them more comprehensive and digestible. Some R or Python packages, such as topGO^10^, GOSemsim^11^ and GOMCL^12^, remove the redundancy of enriched terms by measuring similarities between GO terms. But their results are static table or figures without interactive visualization making them not easy to use for biologists with few programming skills. Some web-based tools were also developed. For example, REVIGO^13^ enables to pre-filter the enriched terms using a cluster algorithm of semantic similarity to find a representative subset and visualize the clustering results with a network map. But it did not include the module to perform functional ORA or GSEA analysis. Another popular tool, Metascape^14^, pre-clustered the enriched terms by Kappa score, and selected the term with the smallest P-value in each cluster as the representative one. However, Metascape only supports ORA analysis and the figures generated by Metascape are non-interactive and static. Users cannot interactively explore the enrichment results. Thus, there are few suitable tools for a comprehensive and user-friendly analysis containing enrichment analysis, redundancy removal, and representative terms selection.

Here, we propose a biologist-oriented web server, named EnrichMiner. It implements two ways with rich interactive operations to visualize the relationships between the terms, reduce the redundancy and select the representatives. One is constructing a network according to pairwise term similarity, clustering the terms with network community detection method. The other is visualizing t the parent-child relationships of the enriched terms through directed acyclic graph (DAG). Additionally, both similarity network map and DAG provides interactive operations to overview the relationships between all the terms, then dive into the relationships between sub terms (in a cluster for pair-wise similarity) and two terms in a pair, and finally view the detail of one term. Along with the maps, there are detailed tutorial videos to help users. EnrichMiner provides interactive operations with the modules in ExpressVis^15^ we developed before. More importantly, EnrichMiner contains a complete pipeline for enrichment analysis. It supports both ORA and GSEA. After uploading a gene list or a gene table with ranked metrics (such as the differential analysis results), users can perform ORA or GSEA, explore the relationships of the enriched terms, select the representatives, and finally visualize them in 7 kinds of downloadable figures. We present two re-analyzed case studies to demonstrate the features and capabilities of EnrichMiner.

## 2 Results

### 2.1 Overview of EnrichMiner

EnrichMiner is a complete pipeline for mining biological insight from omics data. As shown in Fig. 1, its workflow can be divided into three steps: (1) statistical calculation, (2) selection of representative terms after exploring terms’ relationships and diving into the details of one term, (3) visualization of the representative terms.

**Figure 1.**
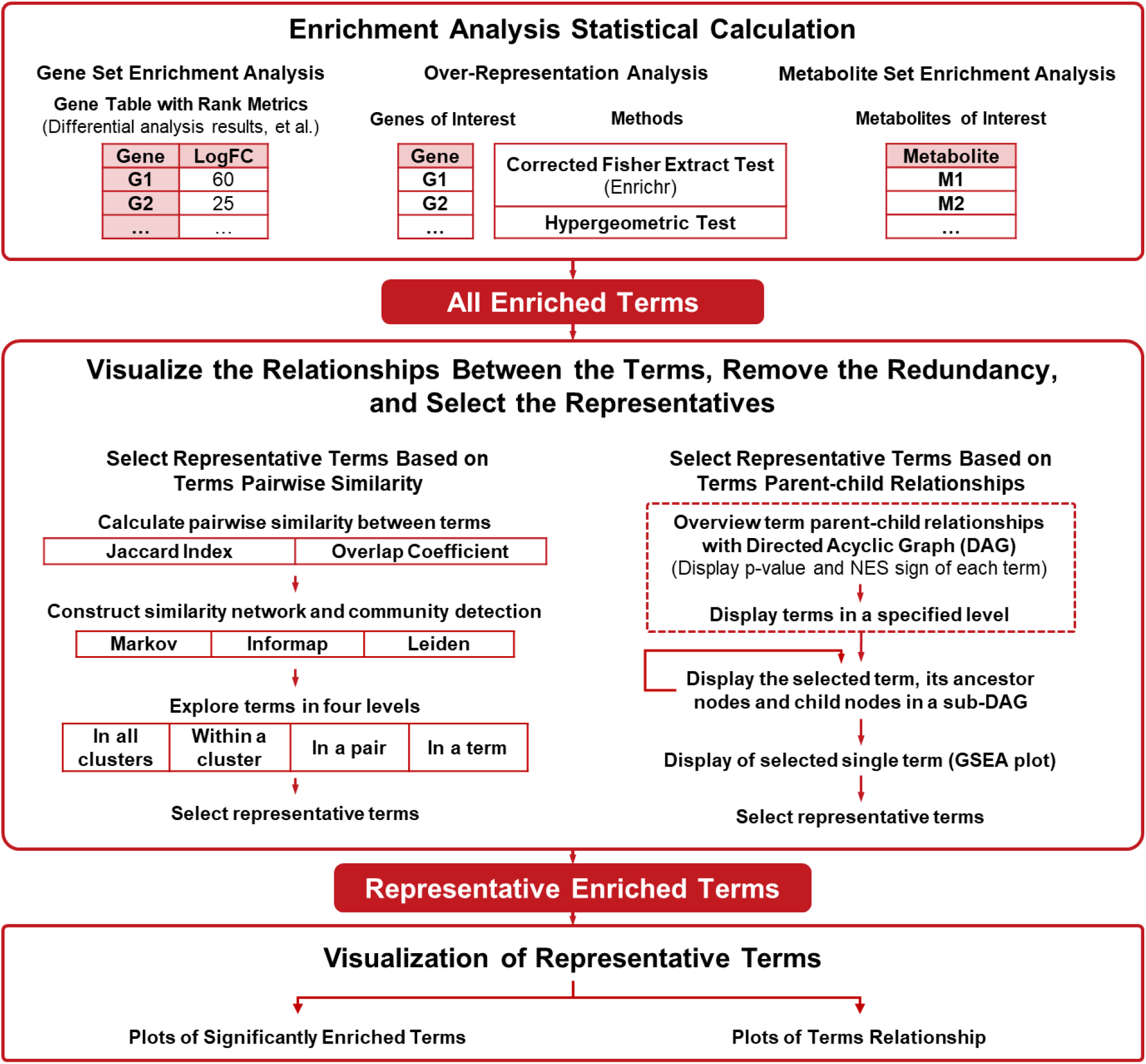
The overview of EnrichMiner workflow. Users can upload a gene table with ranked metrics (for GSEA) or a gene list (for ORA) to calculate the statistical significance of all gene set terms. The most representative terms can be selected with two redundancy reduction methods: community detection of similarity network and term relationship exploration using DAG. Rich interactive operations are available to explore representative terms and relationships among them.

In the statistical calculation step, EnrichMiner provides multiple options for users. At the method level, it supports ORA analysis and GSEA analysis (including MSEA for metabolomics data). ORA takes a gene list as input and GSEA takes a table with ranked metrics as input. At gene set library level, it supports rich gene set libraries that are stored in the database and those uploaded by the users (see Methods for details). At data level, it supports transcriptomics, proteomics, and metabolomics data.

Selection of representative terms is one of the key features of EnrichMiner. It aims to liberate researchers from the heavy manual comparison and empirical selection of enriched terms by visualizing two types of terms’ relationships: pairwise similarity relationship and parent-child relationship. For pairwise similarity relationship, a network based on the pairwise similarity between terms is constructed and the terms are clustered into groups. The network is visualized as a network map. For parent-child relationship, they are visualized as a DAG. Rich interactive visualization operations are provided for users to explore the relationships.

### 2.2 Reduce redundancy and select representative terms

In most commonly-used gene set libraries (like Gene Ontology), many terms contain overlapping genes. The high redundancy of the terms hampers biological interpretations of the enrichment results. In EnrichMiner, to help users explore the terms’ relationships and select representative gene set terms, we implement two methods: one is based on terms pairwise similarity relationship (Fig.2) and the other is based on parent-child relationship (Fig. 3).

**Figure 2.**
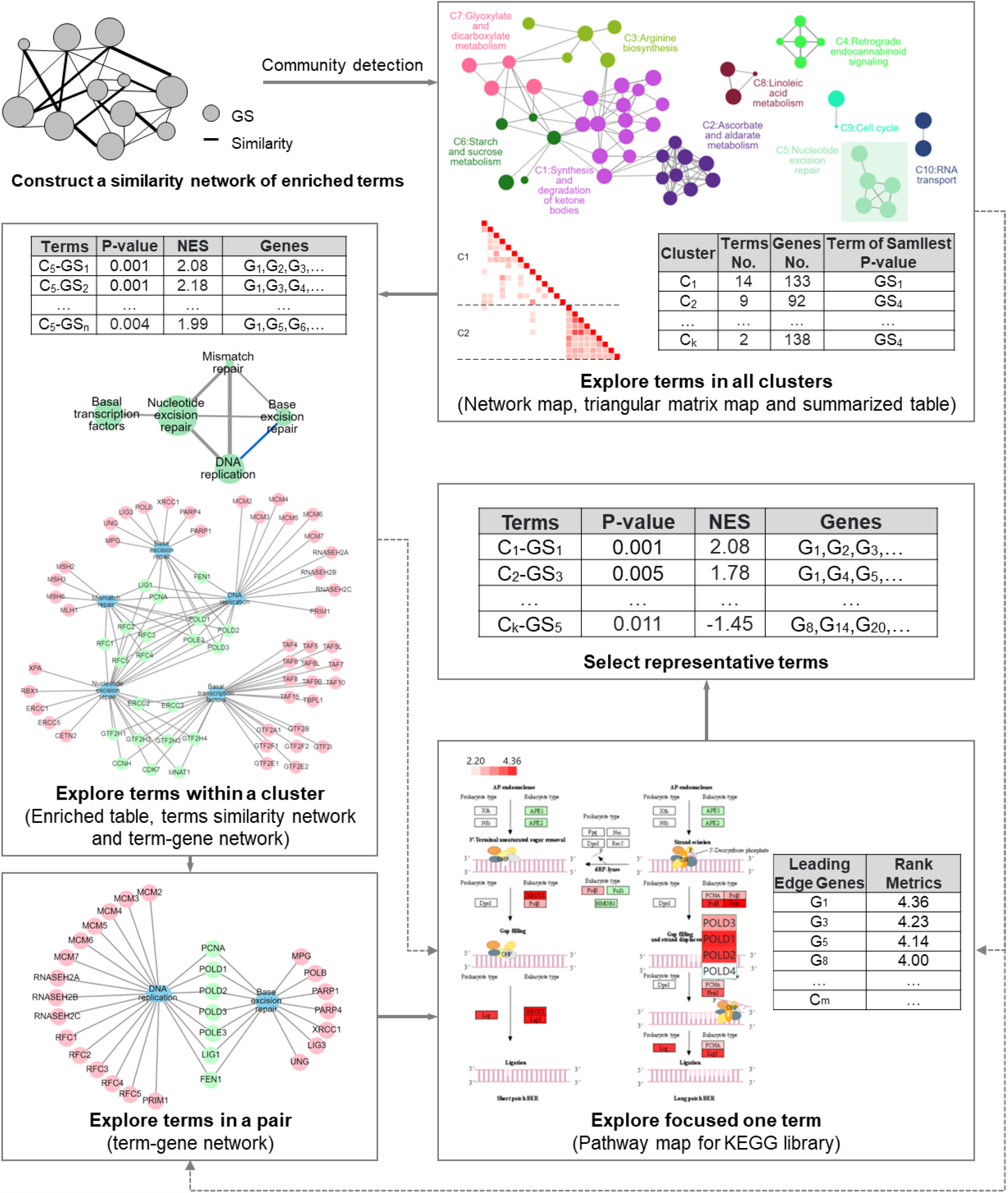
Reduce Redundancy and select representative terms based on terms pairwise similarity. EnrichMiner use enriched terms and their pairwise similarities to construct similarity network, in which nodes represent terms and edges represent the similarity metrics of the corresponding two nodes. Only similarities more than the similarity cutoff are used to construct network. The terms (nodes) are divided into clusters using community detection algorithms. Users can explore the terms in four levels: terms in all clusters, terms within a cluster, terms in a pair and one focused term. Terms similarity network, term-gene network, triangular matrix map, pathway map and various summarized tables are used in these exploration processes.

**Figure 3.**
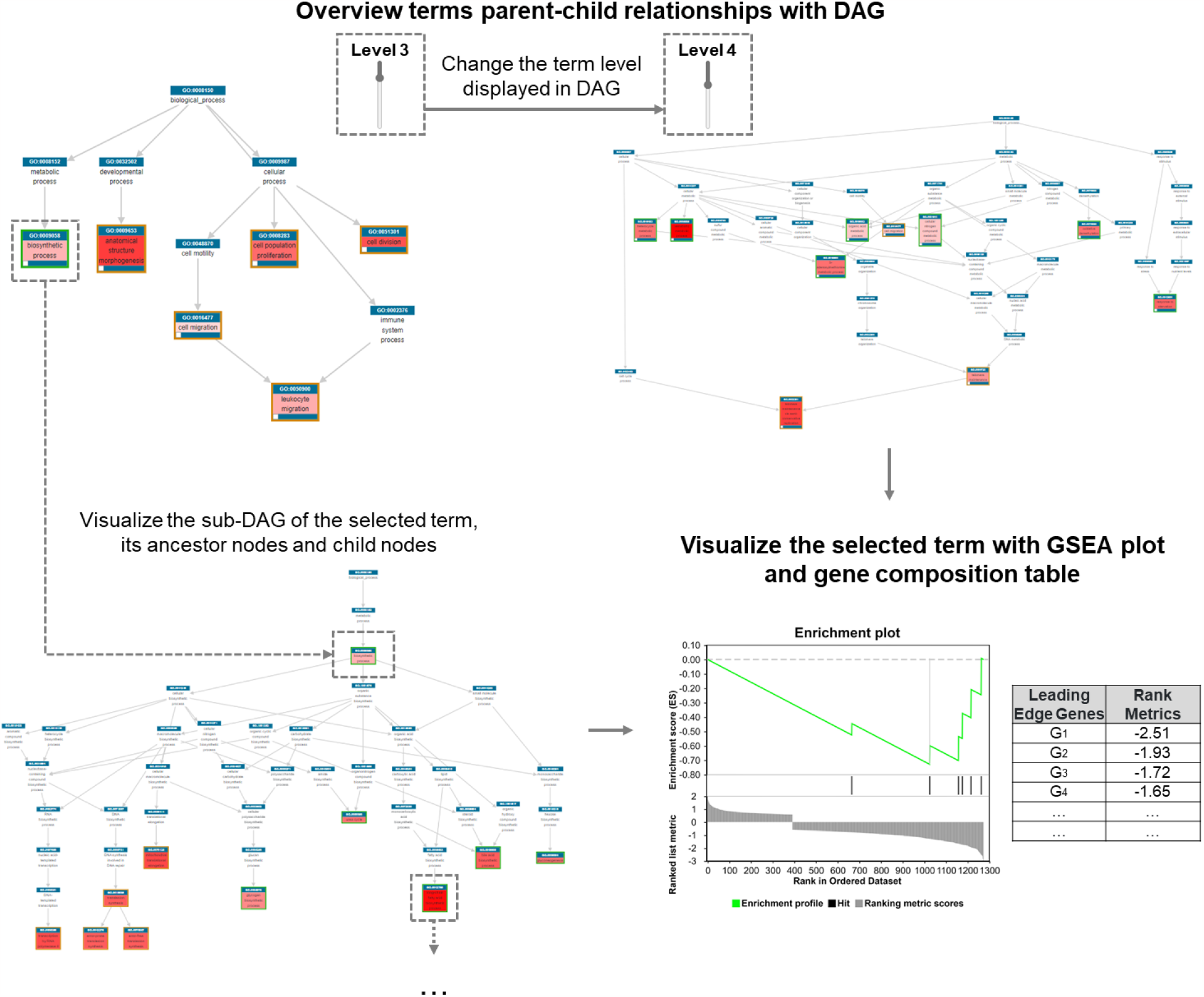
Reduce Redundancy and select representative terms based on terms parent-child relationships. EnrichMiner divides enriched terms into different levels based on the shortest path between the terms and the root term in DAG. Users can decrease or increase the level to overview more general or more specific terms. When users focus on a specific term, they can click on the middle rectangle box of corresponding node to display a sub-DAG of its related ancestor nodes and child nodes.

#### 2.2.1 Reduce redundancy and select representative terms based on terms pairwise similarity

EnrichMiner constructs a similarity network of the enriched terms, where nodes represent terms and edges represent that the similarity between the nodes is more than the similarity cutoff. Then, the nodes (the enriched terms) are clustered into groups through a community detection method (Markov clustering, Informap method or Leiden method). Users can interactively select the similarity calculation method (Jaccard index method or overlap method) and set the similarity cutoff to construct the similarity network, and choose the community detection algorithm to cluster the terms.

To help users explore terms pairwise similarity relationships, EnrichMiner provides interfaces and interactive operations to visualize terms in four levels: terms in all clusters, terms within a cluster, terms in a pair and one focused term. The interfaces of high level provide operations to navigate to the interfaces of low level. Users can overview the results first and then dive into detail in need.

Terms in all clusters are visualized in a network map, a triangular matrix map and summarized table. In the network map, terms with the smallest p-value within each group are selected as represented term and they are displayed as group nodes. Term nodes are displayed with different colors according to the groups they belong to. Node size represents gene number in the term and edge width represents similarity (Fig. S1B). Although the network map made by EnrichMiner can be downloaded for publishing with little modification, users can utilize the following operations to modify the network map: (1) Click to refresh the layout of the network; (2) Move the terms in a group simultaneously; (3) Change or move group label; (4) Change group color (the nodes color changes accordingly); (5) Change the attributes mapped to node size and edge width. In the triangular matrix map, the color represents similarity of the term pairs. The summarized table lists cluster information, such as number of terms, the term with the largest number and the term with the smallest p-value within each cluster.

The network map, triangular map and summarized table provides operations for users to navigate to the interfaces for exploring the terms within a cluster. Three interfaces are provided: a table, a term similarity network and a term-gene network. The table list contains enrichment information, such as p-value, genes in the term. The terms similarity network map displays terms similarity, in which node represents term, edge represents similarity. The term-gene network map displays terms and genes, in which one type of node represents term, another type of node represents genes, the edge between gene node and term node represent that the gene is in the term.

The interfaces for terms in all clusters and those for terms in a cluster provides operations to navigate to the interface for exploring terms in a pair. Two interfaces are provided: a table and term-gene network. The term-gene network map displays terms and genes, in which one type of node represents term, another type of node represents genes, the edge between gene node and term node represent that the gene is in the term. Gene nodes are highlighted with three colors: one for genes in a cluster, one for gene in another cluster and another for genes both in two clusters. Users can easily check the common genes in two terms.

For the term of interest, EnrichMiner provides different interfaces according to gene set library and enrichment analysis method. For the term from the GSEA results, EnrichMiner provides a standard plot modified from raw GSEA software and a table to display the genes information. For the focused term from KEGG pathways, EnrichMiner provides an interface to visualize its pathway map. In the pathway map, the leading-edge genes from GSEA results are highlighted as red color (NES > 0) and blue (NES <0) and the colors are changed according to the values used for ranking; the genes from the ORA results are highlighted as red.

#### 2.2.2 Reduce redundancy and select representative terms based on terms parent-child relationships

Gene sets in some gene set libraries, such as gene ontology and Reactome pathways, have parent-child relationships. They can be represented as directed acyclic graph (DAG) and are arranged hierarchically in different levels from the top term. The low-level (close to the top term) terms in DAG are specific while the high-level ones are general. For the number of the enriched terms can reach tens of, evenly hundreds of, to decrease users’ cognitive burden, EnrichMiner provides level-centered interfaces and operations for users to explore terms relationships.

Initially, EnrichMiner displays the DAG map of the terms under lever 4 and their ancestors. Users can overview the broad terms the genes are enriched in. If they want to display the more specific ones, they can just increase the level. If users focus on one term, they can double the term name to visualize the DAG of the term and its related terms—namely its ancestors and children. For example, if one is interested in metabolism, he/she may want to know which metabolite’s metabolism are enriched in to top ranked genes, while which metabolite’s metabolism are enriched in the low ranked genes (for GSEA). He/she can just click the term metabolism to display its children. This process can be repeated in indefinite times.

In the DAGs mentioned above, EnrichMiner provides three modes: one contains the enriched terms and their ancestors, one contains the enriched terms and the ancestors in the shortest path between them and the root, and another only contains the enriched terms. Users switch between the three modes to visualize according to the number of enriched terms and their needs.

In the DAG map, EnrichMiner provides rich operations to explore the terms: users can drag, move whose DAG map or one DAG node, search and locate terms of their interest. To make DAG map more interactive, the term box is divided into four parts (Fig. S1C): (1) the outer border. (2) the top rectangle box, (3) the middle rectangle box, (4) the lower left check-box. The color of outer border indicates NES of GSEA results (NES < 0, green; NES > 0, brown). The top rectangle box displays the term ID and users can click it to visualize corresponding GSEA plot and gene composition list in GSEA module. The middle rectangle displays the term name, and its background color indicates the enrichment significance (i.e., the darker red indicates the smaller P-values). Sub-DAG related to current node will be shown when users double click this area.

Through the above interactive operations, users can get an overview of the enriched terms and their relationships, and finally determine which terms can be selected as representative terms. Users can check lower left box to select its corresponding term as a representative term and add it into the candidate table.

### 2.3 Metabolite Set Enrichment Analysis

We implemented a light-weighted module for metabolite set enrichment analysis. Compared with the existing tools, the key feature of this module is that users can highlight metabolites on KEGG pathway map or SMPDB pathway maps (Figure S2).

### 2.3 Visualization of the representative terms

The visualization of the representative terms is composed of two types: those display enriched significances and those display relationship among terms (Fig. 4). In both GSEA and ORA modules, interactive scatter plot, bar plot and lollipop plot are generated to show - Log^10^(P-value) of selected terms (Fig. 4A). Additionally, in GSEA module, multi-terms GSEA plot and Rank list of GSEA table are generated to show the normalized enriched score (NES).

**Figure 4.**
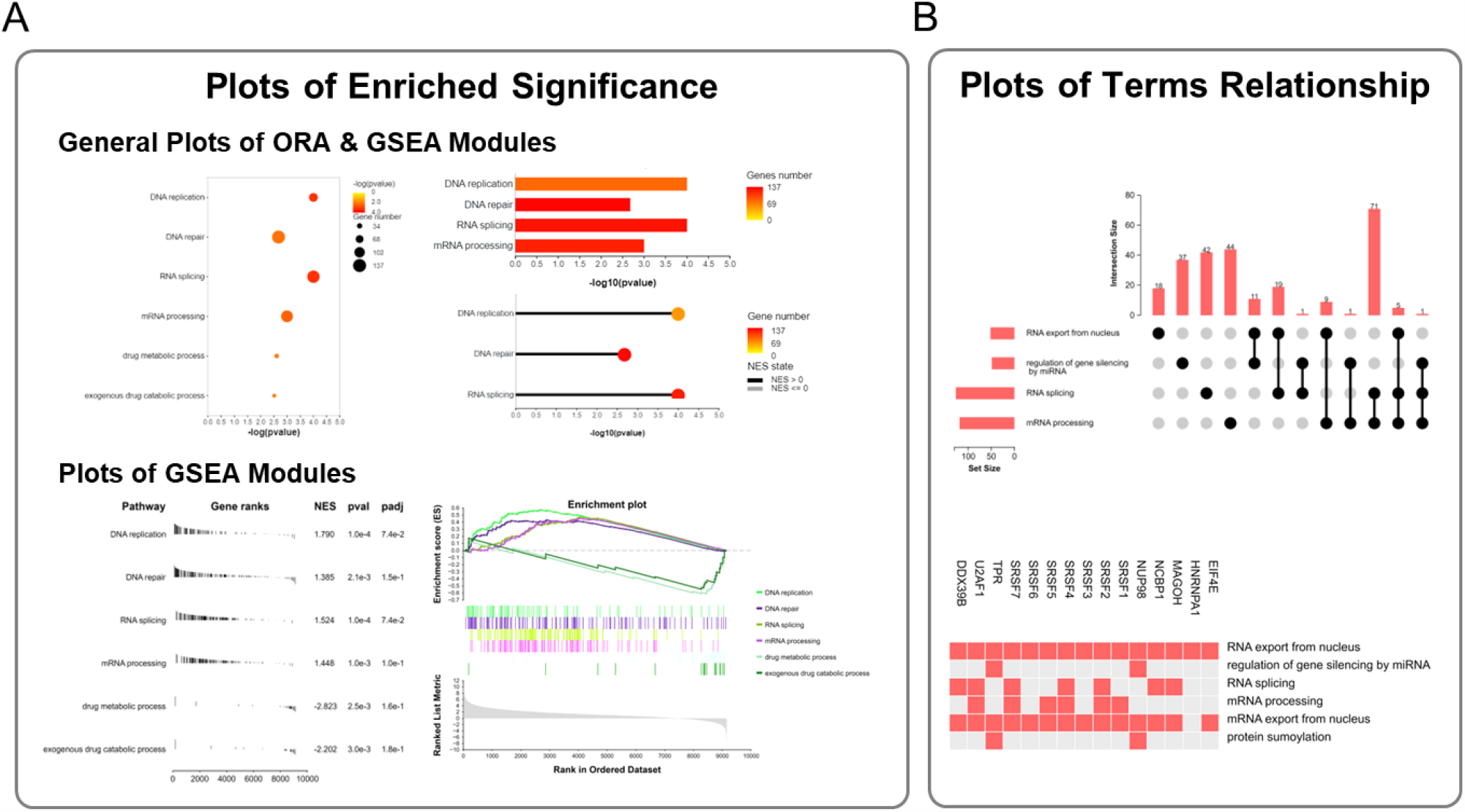
Visualization of EnrichMiner. (A) The statistical significances of enriched terms. (B) The relationships among terms.

Term relationships are displayed via upset plot and “gene set-to-gene” heatmap (Fig. 4B). In upset plot, the cardinality of single terms as well as term combinations with more than 3 gene sets are presented. The total size of each set is represented on the left bar plot. Every possible intersection is represented on the bottom plot, and the number in each subset is shown on the top plot. In heatmap, genes included in each term will be filled with red color.

### 2.4 Case study I: Hepatocellular carcinoma (HCC) and adjacent non-tumor tissue

HCC is one of the most common malignant tumors. To obtain changes in pathways associated with tumorigenesis, we re-analyzed a published HBV-related HCC proteomics dataset^16^ which contains 159 pairs of HCCs (T) and adjacent non-tumor tissues (N).

In GSEA module, we used LogFC(T/N) as the gene ranked and Gene Ontology (GO) Biological Process as the backgroud gene set library. The results show that pathways related to RNA processing and DNA repair are enriched in HCC group. In the contrary, pathways related to metabolism (one of the key liver functions), such as fatty acid metabolism process and P450, are enriched in non-tumor tissue group (Fig. 5A). These indicate that tumors ontain proliferative activity while lose liver function activity..

**Figure 5.**
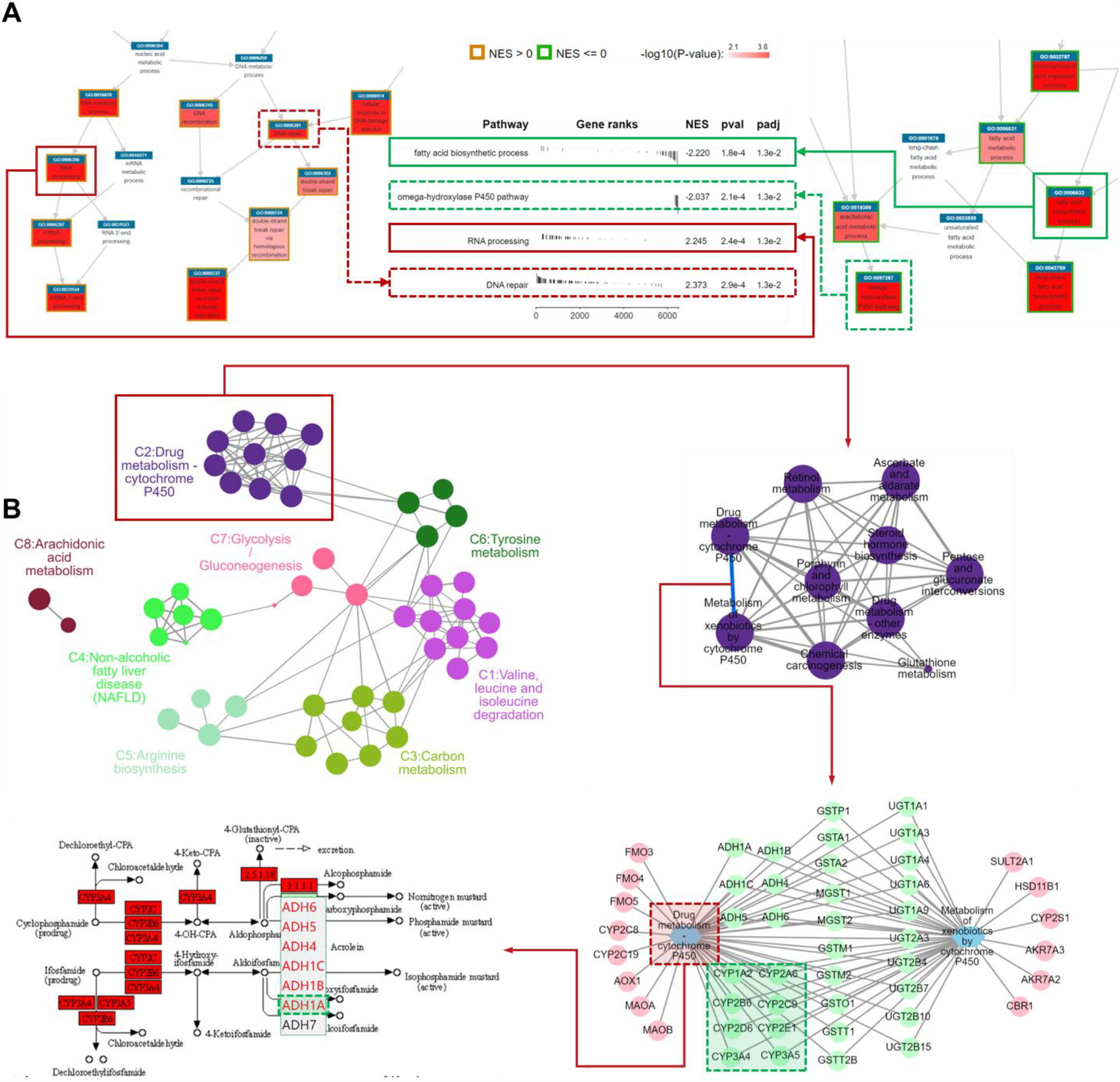
Enrichment results of published HCC Proteomics study (Gao et al.’s cohort). (A) GSEA enriched terms of GO Biological Process gene set database using LogFC(T/N) between tumors and adjacent non-tumors as ranked metrics. (B) ORA enriched terms of KEGG pathway library based on down-regulated proteins between tumors and adjacent non-tumors.

In order to verify the credibility of the above inference that tumors lose liver function activity, we used the down-regulated proteins in tumors (Supplymentary Table 3) for ORA analysis. KEGG library was selected as the background gene set library. The metabolism pathways in liver, such as amino acid degradation, glycolysis/gluconeogenesis, carbon metabolism and drug metabolism, are over-represented (Fig. 5B). We found that multiple Cytochrome P450 (CYP) proteins such as CYP3A, CYP2C, CYP2D, CYP1A and CYP2E are overlapped in multiple drug metabolism-related pathways (green block in term-gene network).

This can be verified by Villeneuve et al.’s and Yan et al.’s studies (drug metabolism is impaired in patients with liver disease^17^ and CYPs are responsible for 70% to 80% of phase I metabolism of 90% of marketed drugs^18^). After highlighting genes on the Drug metabolism - cytochrome P450 pathway map and checking the rate-limiting enzymes (the arrows indicate the reaction in which the enzymes participate are unidirectional rather than bidirectional), we found that ADH1A - an aprognostic biomarker of proteomic subgrouping in the original article^16^ - is among the rate-limiting enzymes.(green block in KEGG pathway map).

### 2.5 Case study II: HCC with and without tumor thrombus

The formation of tumor thrombus is considered a poor clinicopathologic prognostic factor of HCC. To explore the functional biological differences between HCC with and without tumor thrombus, we re-analyzed two published HCC proteome datasets^16, 19^. In Jiang et al.’s cohort, 22 of 110 tumors have tumor thrombus (microscopic vascular invasion-positive, MVI^+^) and in Gao et al.’ cohort, 37 of 159 tumors have tumor thrombus.

We performed differential expression analysis beween HCCs with tumor thrombus and those without tumor thrombus via Welch’s t-test. And the fold change of each feature is calculated based on mean values of samples in each group.

The different Mass spectrometry quantitative methods - label-free DDA for Jiang et al.’s cohort and TMT (Tandem Mass Tag) isotope labeling method for Gao et al.’ cohort - led different qualitative patterns of protein features between two datasets. These two cohorts have 43 common features in the up-regulated proteins and 15 common features in the down-regulated proteins (Fig. 6A).

**Figure 6.**
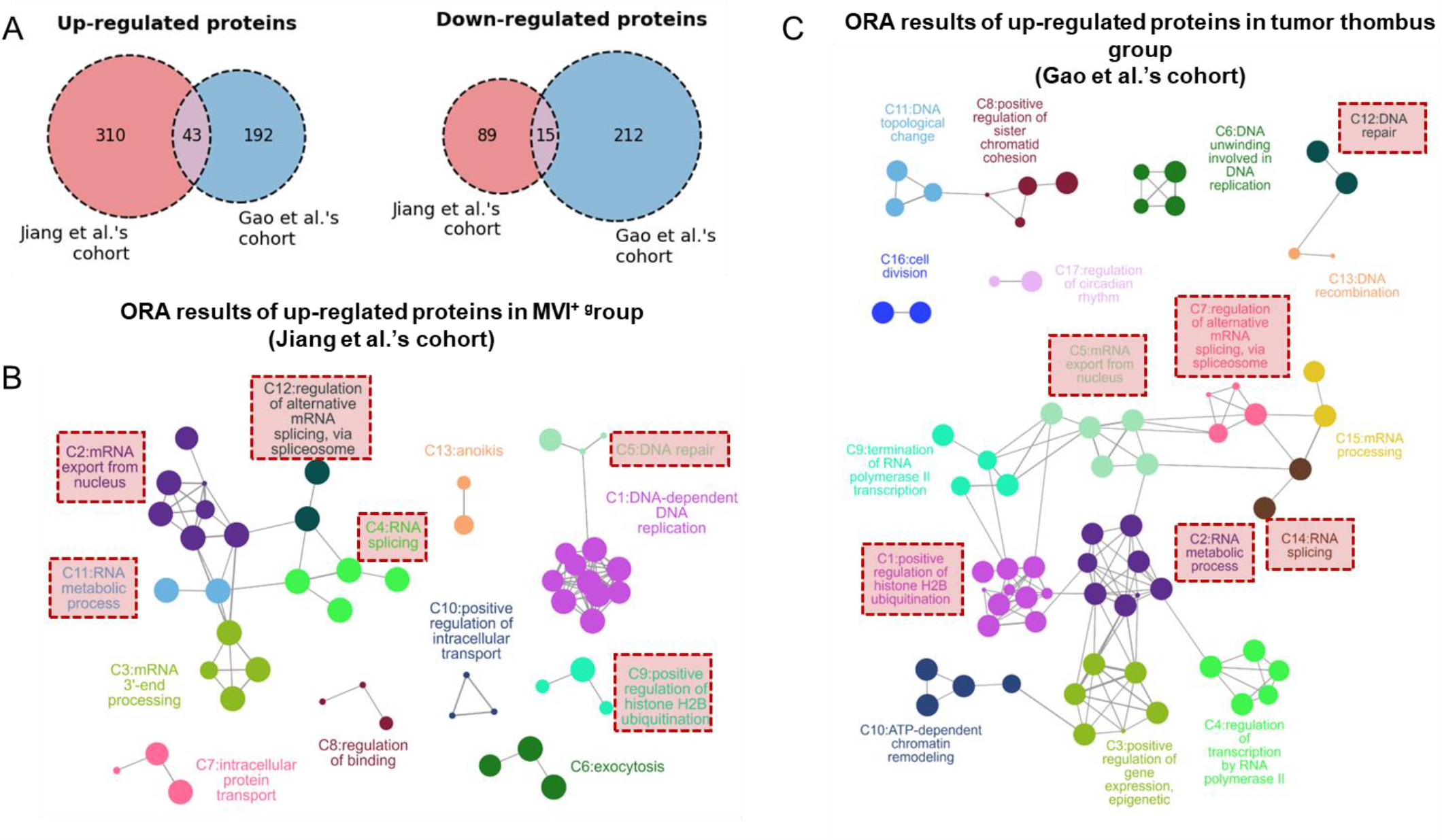
Enrichment results of HCC with or without tumor thrombus based on two published proteomics data sets. (A) Venn diagram of up-/down-regulated proteins between Jiang et al.’s cohort and Gao et al.’s cohort. Pairwise similarity networks based on up-regulated proteins of HCC with tumor thrombus in Jiang et al.’s cohort (B) and Gao et al.’s cohort (C).

We used the down-regulated proteins of each dataset to perform ORA enrichment analysis with Gene Ontology Biological Process library separately, and visualized the relationship between enriched terms through similarity networks (Fig. 6B-C). Unlike the low overlapping at the protein-level, the two datasets enriched similar geneset terms, showing a similar trend in biological functions. HCCs with tumor thrombus are enriched in *mRNA export from nucleus, RNA splicing, RNA metabolic process, DNA repair* and *positive regulation of histone H2B ubiquitination*, which may indicates HCCs with tumor thrombus have more unstable genome and higher transcriptional activity^20^. The results can be confirmed by *regulation of binding* and related terms of *DNA replication* in the Jiang et al.’s cohort and the related terms of *regulation of transcription* in the Gao et al.’s cohort.

## 3 Discussion

Functional enrichment analysis has been used to extract biological insights from -omics data for more than two decades. Although researchers have developed many sophisticated statistical methods, extracting biological insights still requires domain experts’ experience and intuition. To take advantage of experts’ experience and intuition, we should develop tools as a visual analytical platform. EnrichMiner contains well-designed visualization interfaces and rich interactive operations to help users overview the relationships of the enriched terms, explore the relationships of the sub terms, and dive into the details of one term. With EnrichMiner, users can have a better understanding of their functional enrichment results, select representative terms, and generate publishable figures.

In summary, with the well-designed visualization interfaces and rich interactive operations, EnrichMiner is expected to be a useful tool for extracting biological insights from functional enrichment results in an efficient and convenient way. Table 1 lists all the major features of EnrichMiner compared with five other similar web servers. First, EnrichMiner is a complete pipeline, including ORA and GSEA for enrichment analysis, similarity network and DAG for exploring the relationships of the enriched terms and selecting the representatives, and rich interactive and visualization operations for extracting biological insights from representative terms. Second, many user-friendly interactive visualization functions are implemented in EnrichMiner due to the latest web development techniques, and it can be considered as one of the most important features of EnrichMiner.

**Table 1.**
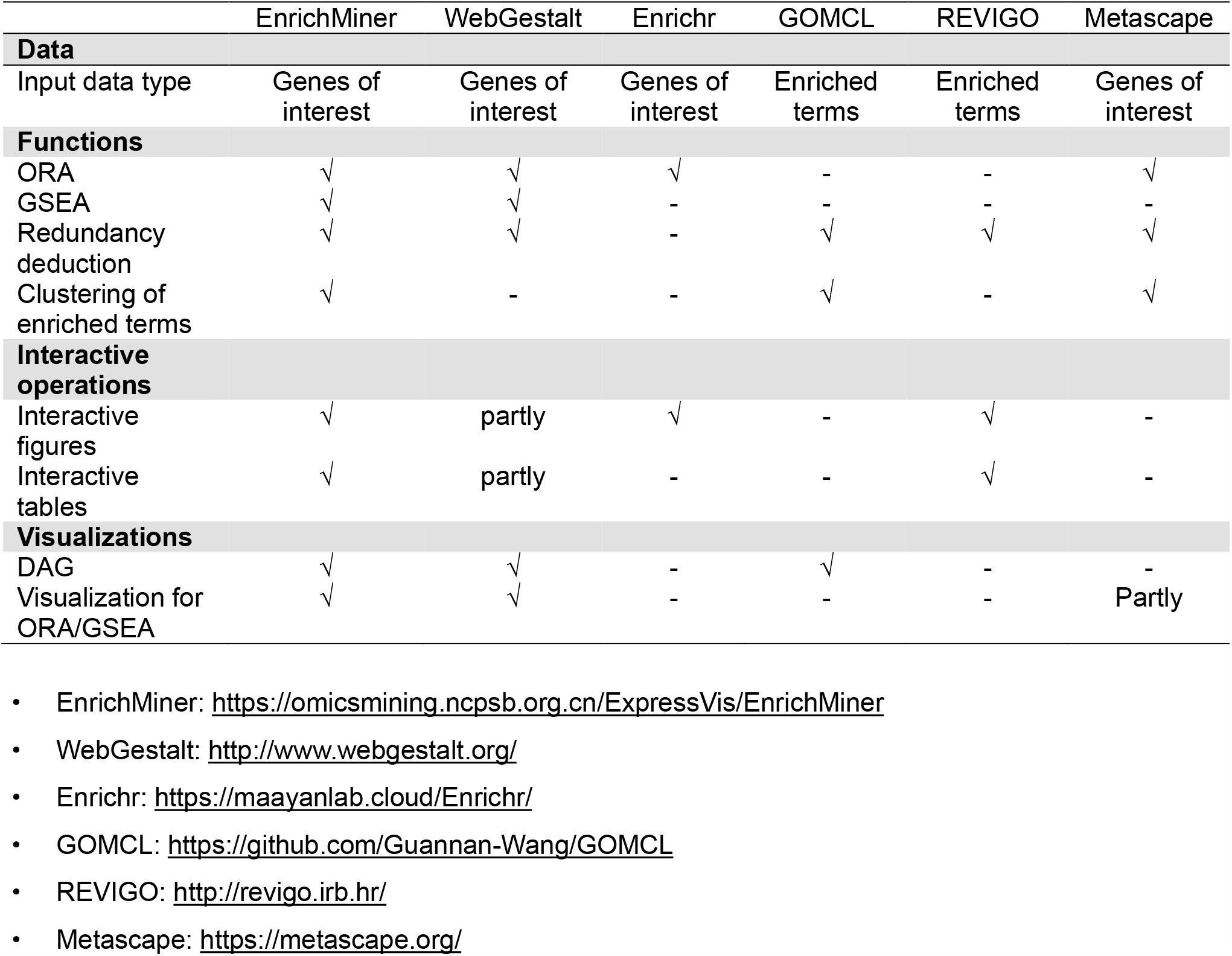
The feature summary of EnrichMiner compared with existing similar tools.

EnrichMiner provides rich interfaces and interactive operations for users to explore the enrichment analysis, and we hope that biologists can participate in the data analysis. However, to better use EnrichMiner, biologists need to learn basic mechanics underlying the statistics and spend some time to control the interactive operations. We have made detailed tutorials. In the future, we should write more intuitive tutorials and introduce EnrichMiner to more biologists.

However, the current version of EnrichMiner has some limitations: the gene set libraries are not enough for the increasing omics studies and an efficient and automatic data synchronization procedure should be considered. In the future, we plan to continue updating EnrichMiner to overcome these limitations. Meanwhile, as large language models (LLM, such as ChatGPT) are keep improving, the way of human-computer interaction will change, and the development of corresponding software tools may experience a paradigm shift. Microsoft is bringing ChatGPT technology to the Office software (such as Word, Excel and Outlook). In the future, we will also pay attention to the integration between EnrichMiner and LLM for better interaction and better user experience.

## 4 Methods

### 4.1 Webserver Architecture

The architecture is similar with that of our previous work ExpressVis. The font-end focuses on visualization and human-computer interaction, the backend focuses on computation. The front-end is implemented with Angular and the server is implemented with Django. EnrichMiner requests Enrichr’s results directly from the browser.

### 4.2 Background Gene Set Library

EnrichMiner includes multiple gene set libraries of model organisms, such as *Homo sapiens, Mus musculus* and *Rattus norvegicus*. The gene set libraries are three sources: MSigDB, Enrichr and those we collected from the gene set databases directly. Additionally, EnrichMiner supports user-defined gene set libraries in the “gmt” format.

For metabolomics, EnrichMiner supports *KEGG compounds* (https://www.genome.jp/kegg) and *SMPDB Primary* (https://smpdb.ca) two metabolite set libraries, which contain 81 and 104 terms respectively. Due to the inconsistency of metabolites nomenclature in published metabolite libraries, we recommend using InChI Key, CAS or KEGG ID of metabolites as input, or executing name matching via our in-home function *Metabolite Name Fuzzy Matching*.

### 4.3 Enrichment Analysis Statistical Calculation

EnrichMiner supports the two mainstream enrichment analysis methods: ORA and GSEA. For ORA analysis, we use hypergeometric test. Raw p-values are adjusted via Benjamini-Hochberg method. The codes are based on Python *scipy* and *statsmodels* packages. In addition, EnrichMiner support ORA analysis provided by Enrichr (https://maayanlab.cloud/Enrichr). For GSEA, we use R package fgsea. Enrichment of metabolomics is also based on ORA method, while only compounds of the chosen metabolite set library are used as background in calculation process.

### 4.4 Pairwise Similarity calculation, network construction and community detection

For each gene set library, we firstly pre-computed the pairwise similarities using python packages *numpy* and *scipy*. The clustering of the network is realized through the community detection algorithms, including Markov clustering, Informap method or Leiden method. Community detection function is implemented with python packages *igraph-python* and *markov_clustering*. Similarity network map is visualized with Cytoscape.js.

### 4.5 Directed acyclic graph

GO obo file is downloaded from gene ontology database. GOATOOLS is used to extract DAG of the terms of interest. DAG is visualized with Cytoscape.js.

### 4.6 Metabolite name matching

EnrichMiner supports exact matching and fuzzy matching of metabolite names. Exact match converts original names to standard names in HMDB database based on corresponding InChI Key or CAS. Fuzzy matching attempts to match the original name with a standard name whose synonyms has a similar Jaro similarity score, which is implemented using the *rapidfuzz* package.

### 4.7 Data availability

Gene lists and gene table referenced in this study are available in Supplementary Data.

### 4.8 Code availability

Custom code used in this study is available from the corresponding authors upon reasonable request.

## Acknowledgment

This work has been supported by the National Key Research and Development Program of China (2021YFA1301604, 2021YFA1301603 and 2020YFE0202200), the National Natural Science Foundation of China (82100130, 32088101), and the CAMS Innovation Fund for Medical Sciences (CIFMS) (2019-I2M-5-063).

## Author Contributions

L.X., XC.B and C.C. designed and co-supervised this project. L.X. and X.T. developed the frontend; L.X. and KK.X. developed the backend. C.C., XC.B, L.X. and KK.X. wrote the manuscript.

## Supplementary Information

**Figure S1.**
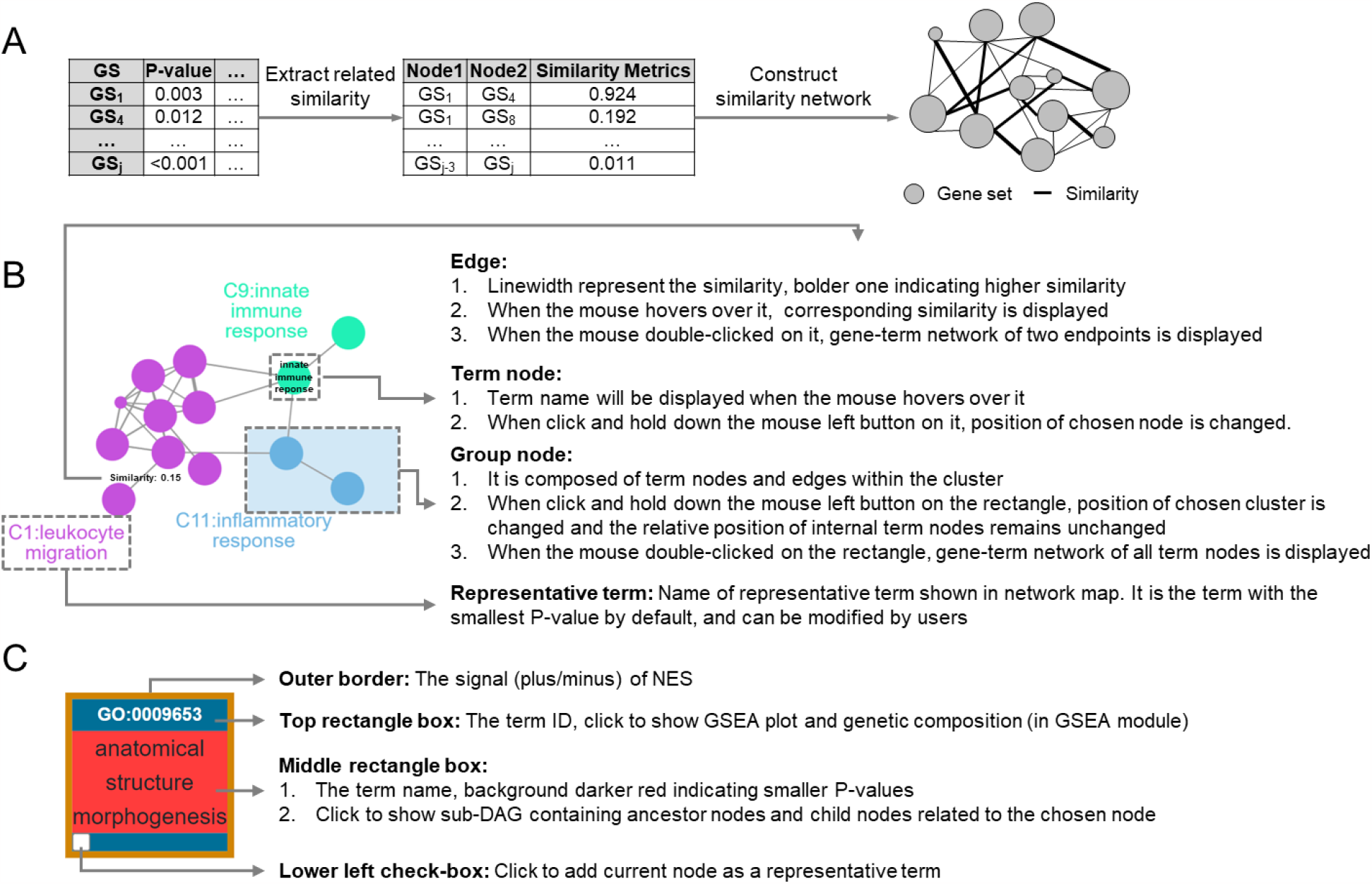
Details of reducing redundancy and selecting representative terms. (A) Processing to construct similarity network. For each gene set library, a data matrix of pairwise similarity is pre-calculated and saved as three columns (Node1, Node2 and Similarity Metrics). In each enrichment analysis task, similarities related to significantly enriched terms are extracted and used to construct similarity network. Nodes represent gene sets and edges represent similarity metrics. (B) Description of nodes and edges in network map. (C) Descriptions of five part in a DAG node (the outer border, the top rectangle box, the middle rectangle box and the lower left check-box).

**Figure S2.**
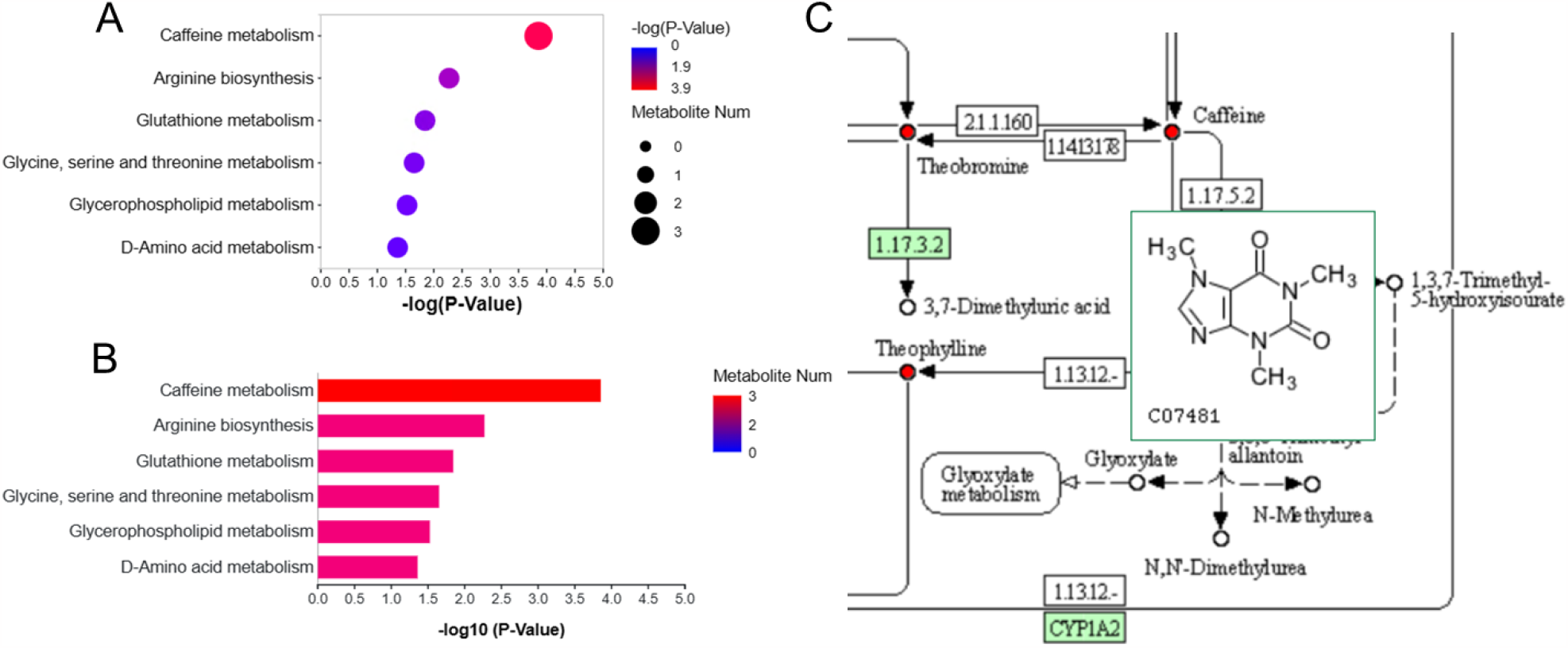
Visualization of metabolite set enrichment analysis. The statistical significances of enriched terms can be displayed with scatter plot (A) and bar plot (B). Metabolites in pathway map can be highlighted and annotated with structural information (C).

